# Exaggerated NMDA Receptor–Primed Metaplasticity via SK Channel Dysregulation in *Fmr1* Knockout Mice

**DOI:** 10.1101/2025.08.26.672454

**Authors:** Toshihiro Nomura, Chad Morton, Anis Contractor

## Abstract

Fragile X syndrome (FXS), the most common monogenic neurodevelopmental disorder associated with autism and intellectual disability, results from the loss of expression of the *FMR1* gene. Synaptic and circuit-level abnormalities are well documented in FXS and extensively studied in the *Fmr1* KO mouse model. In CA1 hippocampal neurons functional, molecular and structural synaptic changes have been described yet the canonical form of Hebbian CA1 long term potentiation (LTP) remains intact in *Fmr1* KO mice. Here we examined whether state-dependent synaptic plasticity in CA1, in which prior “priming” activity modulates subsequent synaptic plasticity, was affected in *Fmr1* KO mice. We found that NMDA receptor activation prior to LTP induction produced metaplastic inhibition of LTP, which was exaggerated in *Fmr1* KO mice. This effect was mediated by the activity of small conductance calcium-activated potassium (SK) channels which was enhanced after NMDA priming, and dampened dendritic excitability. Blocking SK channels during NMDA-primed LTP induction eliminated the abnormal metaplasticity in *Fmr1* KO slices, implicating altered SK activity in the exaggerated LTP inhibition in *Fmr1* KO mice. These finding reveal a disrupted coupling between NMDA receptors and SK channels in *Fmr1* KO mice, which alters the impact of priming on LTP expression in the CA1. Altered metaplasticity may represent a neural correlate of impaired adaptive hippocampal learning in *Fmr1* KO mice.

**Significance Statement:** While conventional synaptic plasticity (LTP and LTD) has been extensively examined in *Fmr1* KO mice, evidence about the integrity of metaplasticity in these mice has been limited. This study provides a characterization of alterations in NMDA receptor mediated metaplasticity in the hippocampus in *Fmr1* KO mice. The question of whether hippocampal LTP is altered in these mice remains unresolved, and changes in metaplasticity may partly explain the discrepancies across studies. Our findings not only identify novel synaptic phenotypes and their underlying mechanisms in the FXS mouse model, but also highlight potential therapeutic targets for FXS.

## Introduction

Fragile X syndrome (FXS) is the most common single-gene cause of autism and intellectual disability. It arises from a CGG repeat expansion in the 5’ untranslated region of the *FMR1* gene leading to loss of expression of the fragile X messenger ribonucleoprotein (FMRP) (Hagerman et al., 2017). FMRP is an RNA-binding protein that directly or indirectly regulates the expression of many other genes including many that encode for synaptic proteins (Richter and Zhao, 2021). This has propelled intensive investigations into how synaptic mechanisms, including synaptic plasticity, may be altered by the loss of Fmrp (Booker and Kind, 2021). In this respect, the use of rodent models of the disorder have been central to establishing the cellular, synaptic and circuit pathology of FXS (Bear et al., 2004)(Sidorov et al., 2013)(Salcedo-Arellano et al., 2020).

Synaptic plasticity is critical both to developmental processes that underlie neural circuit formation particularly during critical periods (Reh et al., 2020), as well as to cognitive processes such as learning and memory (Citri and Malenka, 2008). Multiple forms of synaptic plasticity have been described in the brain that regulate the efficacy of synaptic transmission through distinct mechanisms and effectors. The canonical forms of synaptic plasticity have been studied in the mammalian hippocampus because Hebbian plasticity has been proposed as the biological substrate of learning and memory. Synaptic plasticity has been a focus in many neurodevelopmental disorders and their animal models including in the *Fmr1* KO mice (Sidorov et al., 2013). These mice have been demonstrated to have an altered protein synthesis dependent form of long-term depression (LTD) in synapses of CA1 neurons (Huber et al., 2002)(Nosyreva and Huber, 2006). Conversely most initial studies reported no major changes in CA1 long-term potentiation (LTP) in *Fmr1* KO mice (Godfraind et al., 1996)(Paradee et al., 1999)(Li et al., 2002)(Larson et al., 2005)(Bostrom et al., 2015)(Bostrom et al., 2015)(Talbot et al., 2018)(Brager et al., 2012). Later work using threshold induction protocols, revealed impairments in CA1 LTP (Lauterborn et al., 2007; Hu et al., 2008; Lee et al., 2011) while under different experimental conditions a weak theta burst induction that induced backpropagating action potentials resulted in more easily induced LTP in *Fmr1* KO slices (Routh et al., 2013). While these studies appeared somewhat contradictory, the different paradigms utilized for LTP induction suggest that changes in the threshold for the induction of plasticity could be affected by the state-dependence of the neuron.

State dependent plasticity, or metaplasticity, is a higher order regulation of synaptic plasticity in which prior cellular or synaptic activity modifies the capacity for subsequent plasticity, endowing synapses with the ability to integrate plasticity signals over time (Abraham and Bear, 1996). *In vitro*, priming stimuli that do not by themselves cause lasting changes can either promote or suppress the expression of subsequent synaptic plasticity. Several signaling pathways mediate metaplasticity, with N-methyl-D-aspartate (NMDA) receptors (NMDARs) and Group 1 metabotropic glutamate receptors (mGluRs) playing prominent roles in the hippocampus (Abraham, 2008). Priming stimuli that activate NMDA receptors inhibit subsequent synaptic plasticity induction (Huang et al., 1992) while mGluR priming promotes synaptic plasticity (Cohen et al., 1998).

There is evidence that some mechanisms of metaplasticity are altered in *Fmr1* KO mice. For example while mGluR priming enhances subsequent CA1 LTP to the same magnitude, it becomes independent of the requirement for the protein synthesis during induction in the KO mice (Auerbach and Bear, 2010). In contrast, NMDAR priming dependent inhibition of CA1 LTP has not been systematically examined in *Fmr1* KO mice.

In the present study, we tested whether NMDAR-dependent metaplasticity is altered in the CA1 region of the hippocampus in *Fmr1* KO mice. We found NMDAR activation prior to LTP induction significantly inhibited hippocampal LTP in *Fmr1* WT and KO mice. However, when a stronger induction protocol was applied, the inhibitory effect of NMDA priming was overcome in WT mice, but not in *Fmr1* KO slices indicating enhanced priming effects in *Fmr1* KO mice. This exaggerated inhibition was mediated by abnormal modulation of a small conductance Ca^2+^ activated potassium (SK) channel during metaplasticity. These findings demonstrate that dysregulated functional coupling between NMDARs and SK channels in *Fmr1* KO mice alters state-dependent regulation of synaptic potentiation, potentially impairing cognitive processes that rely on prior experience (Abraham, 2008).

## Results

### Priming by NMDA causes greater LTP suppression in Fmr1 KO mice

Prior studies using a strong LTP induction protocol have demonstrated that the magnitude of early and late phase hippocampal CA1 LTP is comparable to WT mice in *Fmr1* KO mice (Godfraind et al., 1996; Paradee et al., 1999; Li et al., 2002; Larson et al., 2005; Bostrom et al., 2015; Talbot et al., 2018). Using a single theta-burst stimulation (TBS) for induction we found that the magnitude of LTP was indistinguishable between *Fmr1* WT and KO mice (WT: 140.4 ± 8.4 %, n = 7 slices, 4 mice; KO: 135.4 ± 10.7 %, n = 7 slices, 4 mice; p = 0.90; Mann-Whitney) (Figure 1A & B). Synaptic metaplasticity and homeostatic mechanisms are altered in *Fmr1* KO mice (Auerbach and Bear, 2010; Chen et al., 2022) but to date there have not been any assessment of one of the characteristic forms of metaplasticity in the CA1 that is triggered by NMDA receptor activation and depresses subsequent LTP expression (Huang et al., 1992; Abraham, 2008; Zorumski and Izumi, 2012). We examined NMDA receptor mediated metaplasticity using pre-conditioning with pharmacological activation of NMDA receptors (1 μM, for 10 min, 5 min prior to LTP induction) (Izumi et al., 1992b; Izumi et al., 1992a; Kato et al., 1999; Izumi et al., 2008). Subsequent TBS LTP was inhibited equally in both *Fmr1* WT and KO mice with no potentiation observed at 50-60 minutes post induction (WT: 105.1 ± 9.1 %, n = 8 slices, 4 mice; KO: 102.3 ± 4.7 %, n = 11 slices, 5 mice; p = 0.97; Mann-Whitney) (Figure 1 C & D).

**Figure 1.**
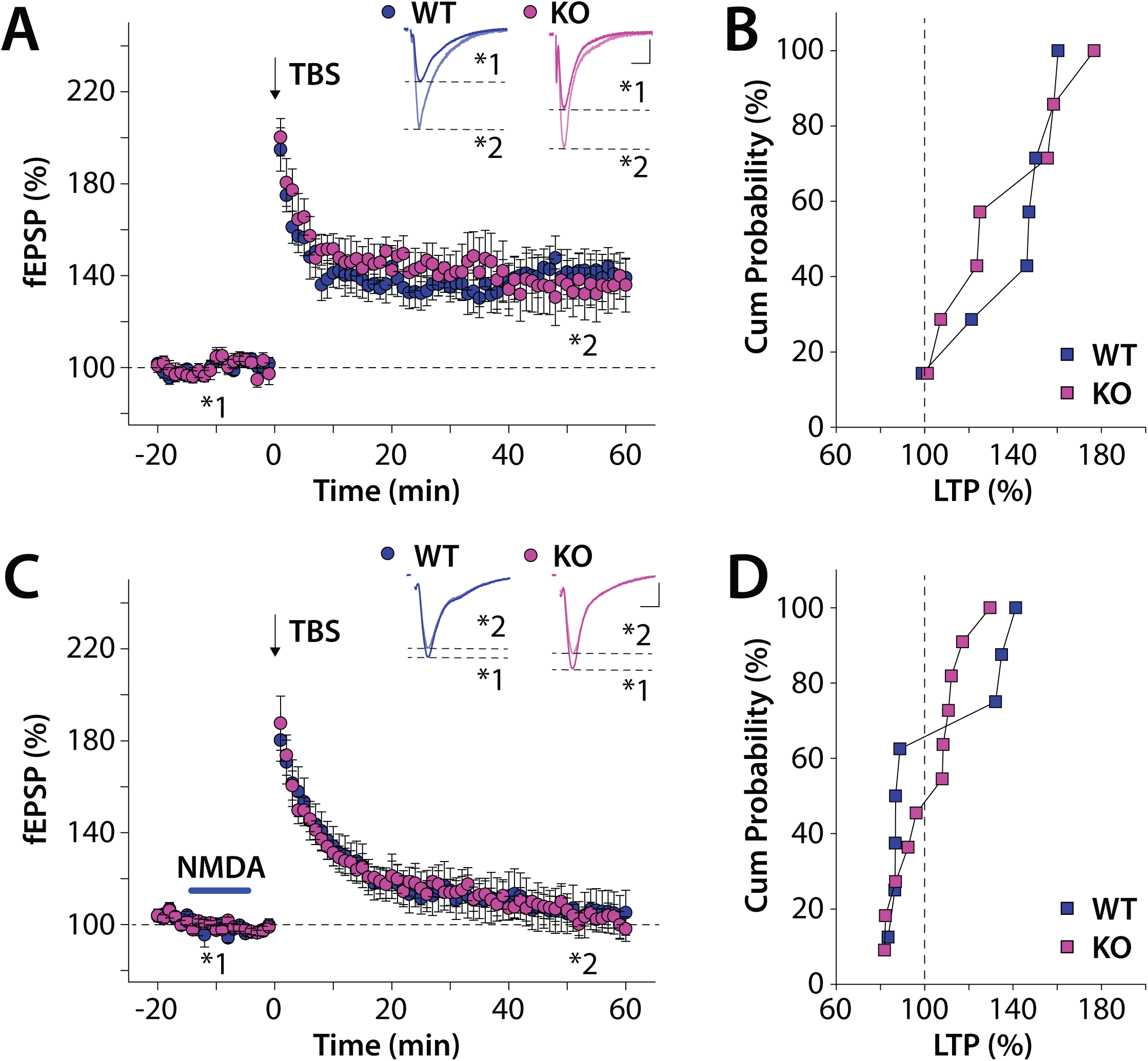
CA1 LTP and NMDA priming induced metaplastic LTP inhibition in *Fmr1* KO mice. **A**) Time course of CA1 LTP induced by theta burst stimulation (TBS; 5 bursts at 100 Hz repeated by 15 times at 5 Hz theta frequency) in *Fmr1* WT and KO mice. Inset shows fEPSP traces before and after LTP from a representative recording. Calibration: 10ms, 0.2 mV. **B**) Cumulative probability graph of potentiation after LTP in each recording. **C**) Time course of TBS induced LTP with NMDA priming (1μM for 10 min). Inset are representative traces of fEPSPs during baseline recording and post LTP induction (50 - 60 min bin after TBS). Calibration: 10ms, 0.2 mV. **D**) Cumulative probability of potentiation after NMDA priming

To determine whether a stronger repeated LTP induction protocol was similarly inhibited by NMDA priming we used a theta burst stimulus repeated 5 times (5x TBS) with an intertrain interval of 2 mins for induction. 5x TBS LTP was comparable between *Fmr1* WT and KO slices (WT: 191.1 ± 21.7 %, n = 10 cells, 5 mice; KO: 181.4 ± 6.5 %, n = 10 cells, 5 mice; p = 0.68; Mann-Whitney) (Figure 2 A & B). When the 5x TBS induction was preceded by a 10-minute priming with low concentration of NMDA, significant potentiation was observed 60 minutes after the last TBS train in slices from WT mice (Figure 2 C & D). In contrast, in slices from *Fmr1* KO mice, in pre-primed slices the 5x TBS induction produced normal post-tetanic potentiation but 60 minutes after the last train no potentiation was measured in most slices, (WT: 126.2 ± 9.1 %, n = 8 cells, 4 mice; KO: 94.3 ± 5.4 %, n = 8 cells, 3 mice; p = 0.024; Mann-Whitney) (Figure 2C & D). These results indicate that NMDA-priming induces aberrant metaplastic LTP suppression in *Fmr1* KO mice.

**Figure 2.**
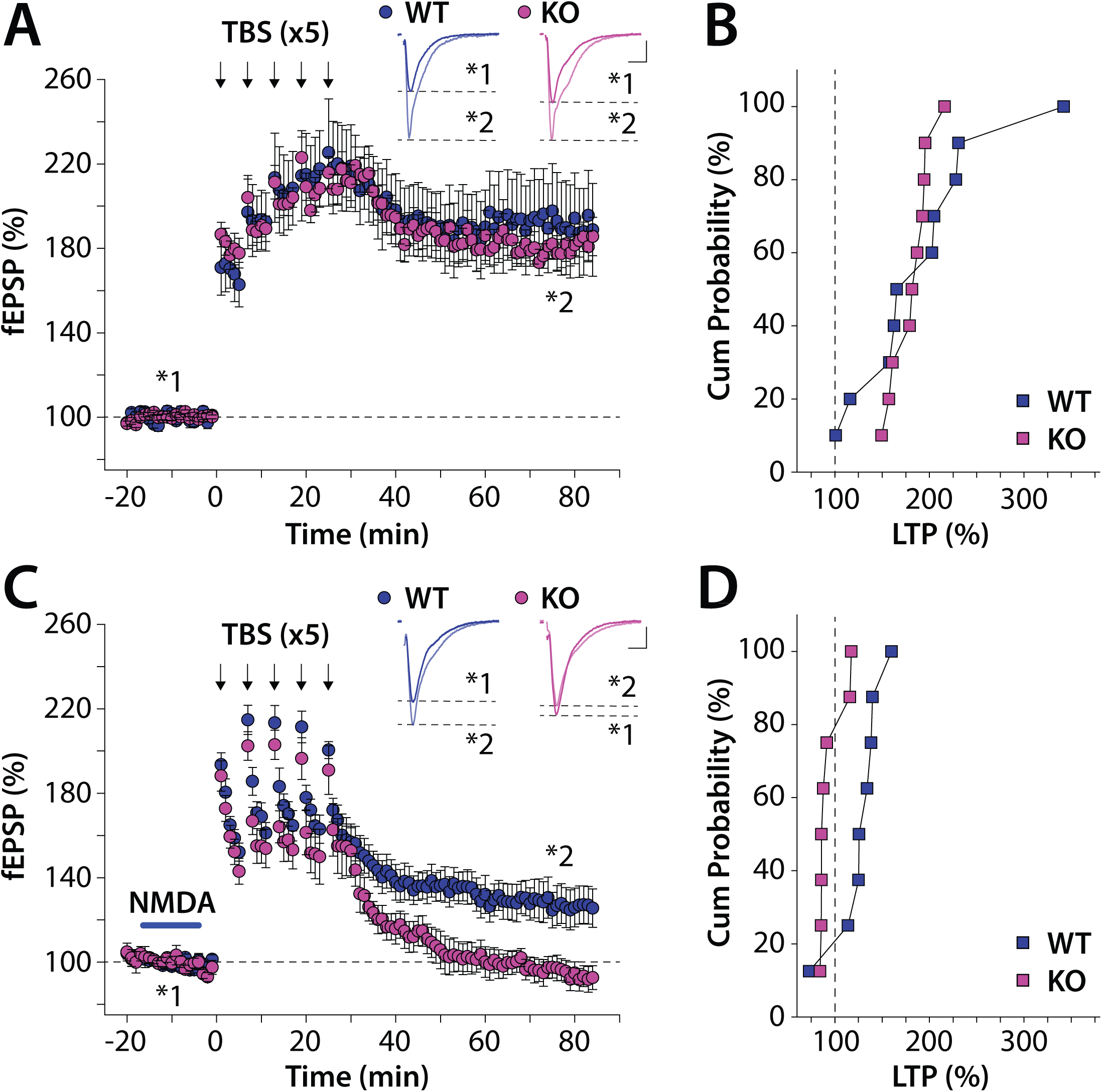
CA1 LTP and metaplasticity after a prolonged TBS induction in *Fmr1* KO mice. **A)** Time course of LTP induced by five repeated TBS stimuli (2 min intervals). Inset shows representative fEPSP traces. Calibration: 10ms, 0.2 mV. **B)** Cumulative probability of potentiation in WT and KO slices after repeated TBS **C**) Time course of NMDA priming metaplasticity with repeated TBS in WT and KO slices. Inset are representative traces of fEPSPs during baseline recording and post LTP induction. Calibration: 10ms, 0.2 mV. **D**) Metaplasticity LTP data for each slice represented as the cumulative probability.

### NMDA priming stimuli suppress dendritic excitability aberrantly in Fmr1 KO mice

The physiological substrates for NMDA priming are not clearly elucidated (Abraham, 2008). As NMDA receptors are important for local effects on the excitability of dendrites (Losonczy and Magee, 2006) which in turn can affect LTP induction (Kim et al., 2007) we hypothesized that NMDA priming might dampen dendritic excitability to cause metaplastic LTP inhibition. To test this, we recorded Ca^2+^ transients in the dendrites of CA1 neurons as a proxy for backpropagating action potentials (bAPs) evoked by somatic depolarization using 2-photon laser scanning microscopy (2PLSM) (Day et al., 2008; Nomura et al., 2023) (Figure 3 A). To determine if NMDA priming affected dendritic excitability, we measured the effect of a 10 mins NMDA treatment on the dendritic Ca^2+^ signal. In recordings from WT slices we found no difference in the Ca^2+^ response along the dendrite before and after NMDA treatment (Figure 3 B & D) (p = 0.20; Wilcoxon). However, in recording from *Fmr1* KO slices there was a significant reduction in the dendritic Ca^2+^ signal following NMDA priming (Figure 3C & E) (p < 0.0001; Wilcoxon). The ratio of the Ca^2+^ signal before and post NMDA application was reduced in *Fmr1* KO mice compared to in WT mice (WT: 90.5 ± 10.0 %, n = 11 cells, 5 mice; KO: 63.1 ± 6.7 %, n = 16 cells, 7 mice; p = 0.028; Kolmogorov–Smirnov) (Figure 3F). These results indicate that there is an aberrant NMDA induced suppression of dendritic excitability in *Fmr1* KO mice that potentially contributes to exaggerated metaplastic LTP inhibition.

**Figure 3.**
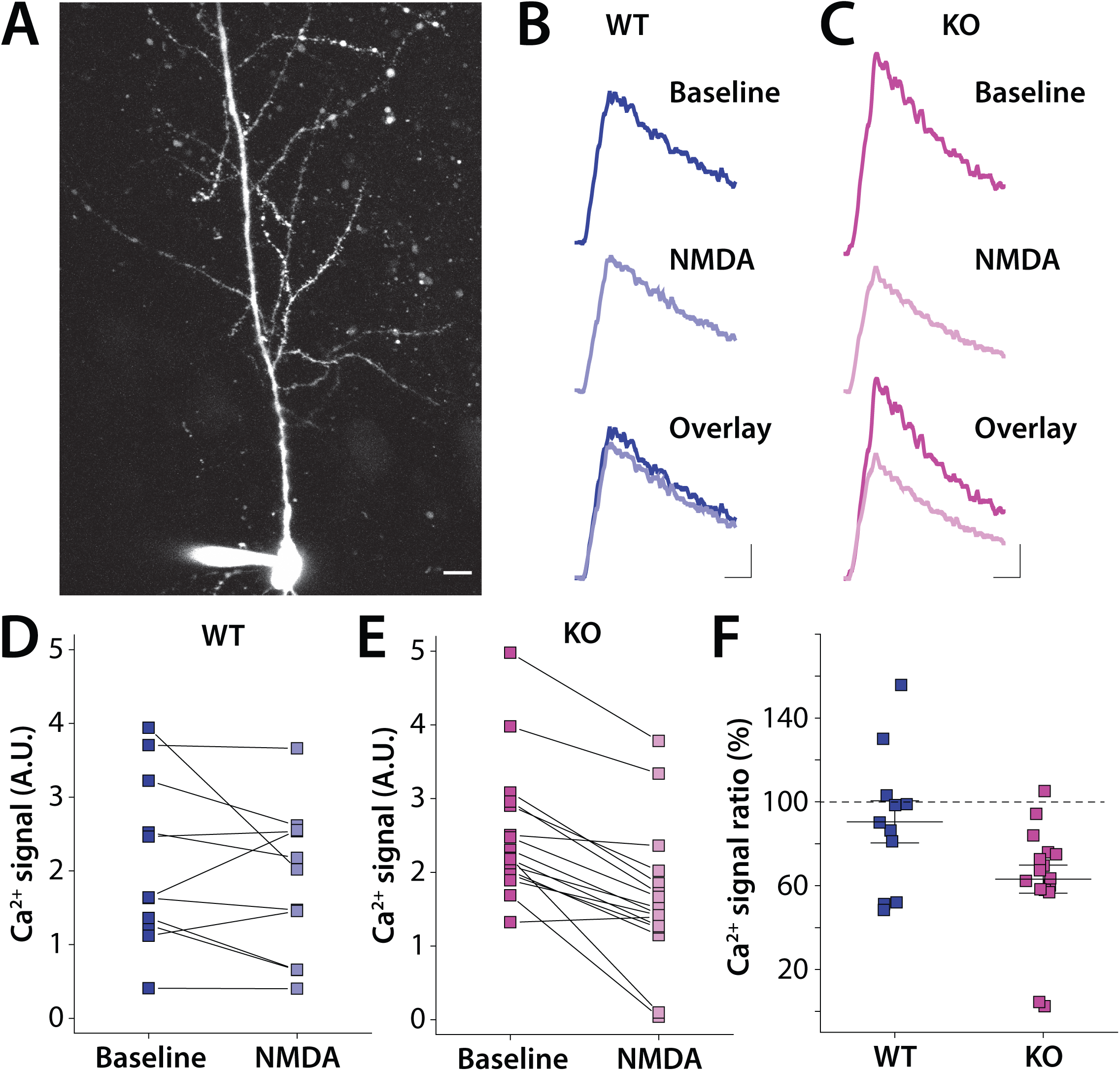
Dendritic excitability is inhibited after NMDA priming in *Fmr1* KO slices. **A)** Example 2PSLM image of CA1 pyramidal neurons. Dendritic Ca^2+^ signals were recorded in ROIs located in apical dendrites ∼100 μm distal from the soma. Ca^2+^ signals were activated by backpropagating action potentials (bAPs) evoked by somatic suprathreshold depolarization. Calibration: 10 μm. **B**) Representative Ca^2+^ transients (Λ1F/F) recorded at baseline and after NMDA priming from *Fmr1* WT mice and **C**) *Fmr1* KO slices. Calibration: 500 ms, 0.5 arbitrary unit. **D**) Grouped data from all recordings of Ca^2+^ signal at baseline and after NMDA priming in WT and **E)** *Fmr1* KO slices. **F**) Grouped data for the change (% of baseline) in Ca^2+^ signal after NMDA priming in WT and KO neurons.

### NMDA receptor activity is not affected in Fmr1 KO mice

NMDA receptors play a critical role in LTP induction, and during priming signaling pathways downstream of NMDA receptors are active (Zorumski and Izumi, 2012). Prior work demonstrated that in young mice (2 weeks) but not older (6 weeks) *Fmr1* KOs NMDA receptor synaptic expression was reduced (Pilpel et al., 2009). Therefore it is possible that in our study in which we use mice between 3-5 weeks direct alterations in NMDA receptors might affect metaplasticity in *Fmr1* KO mice. To examine the contribution of NMDA receptors to synapses in the CA1 we first measured the ratio between the amplitudes of NMDA receptor mediated component (EPSC_NMDA_) and AMPA receptor mediated component (EPSC_AMPA_) of the evoked EPSC. The NMDA/AMPA ratio (EPSC_NMDA_/EPSC_AMPA_) was not significantly different between recordings in slices from *Fmr1* WT and KO mice (WT: 31.1 ± 2.4 %, n = 31 cells, 7 mice; KO: 26.1 ± 2.7 %, n = 37 cells, 9 mice; p = 0.10; Mann-Whitney) (Figure 4 A & B). We found that EPSC_AMPA_ was also indistinguishable between *Fmr1* WT and KO mice as the input-output curves of EPSCs recorded at V_m_ = -70 mV did not diverge between groups (WT: n = 17 cells, 7 mice; KO: n = 20 cells, 6 mice; F_(1,35)_ = 0.04, p = 0.83; two-way ANOVA) (Figure 4 C & D). While the amplitude of EPSC_NMDA_ was not altered, we found that the decay kinetics were significantly slower in *Fmr1* KO mice than in WT mice (WT: 83.5 ± 8.3 ms, n = 9 cells, 4 mice; KO: 133.8 ± 8.6 ms, n = 12 cells, 4 mice; p = 0.002; Mann-Whitney) (Figure 4 E & F). The subunit composition of NMDA receptors is a major factor that determines channel kinetics where GluN2B subunit containing receptors demonstrate a slower time course of decay (Paoletti et al., 2013). In addition, GluN2B containing NMDA receptors play a role in NMDA receptor dependent metaplasticity (Izumi et al., 2008). Previous studies also reported an increase in GluN2B protein level in *Fmr1* KO mice by biochemical analyses (Schutt et al., 2009)(Toft et al., 2016). Therefore, we tested if the EPSC_NMDA_ was differentially affected by the GluN2B selective antagonist ifenprodil (IFN) which would indicate a change in the relative contribution of this subunit in *Fmr1* KO mice. However, we found that IFN (5 μM) inhibited EPSC_NMDA_ similarly in *Fmr1* WT and KO mice (WT: 60.6 ± 4.6 %, n = 9 cells, 4 mice; KO: 57.9 ± 3.5 %, n = 12 cells, 4 mice; p = 0.46; Mann-Whitney) (Figure 4G & H). These results indicate that there are no gross changes detected in the NMDA receptors in CA1 synapses in the ages of mice used in our study that could account for the dysregulated metaplasticity in *Fmr1* KO mice.

**Figure 4.**
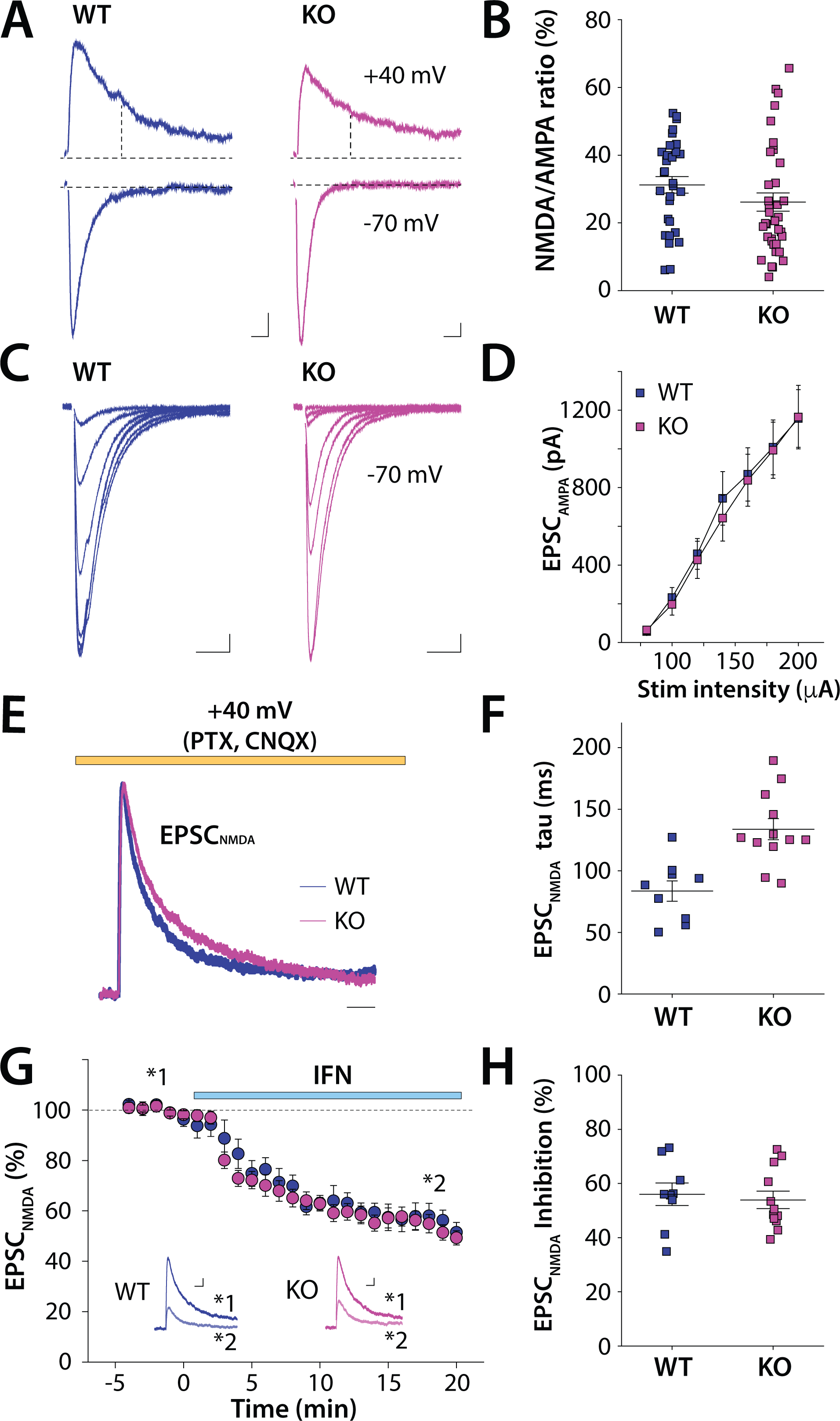
Synaptic AMPA and NMDA receptor mediated EPSCs in CA1 are not disrupted in 3-5 week old *Fmr1* KO mice. A) Representative EPSC traces in CA1 neurons recorded at V_m_ = +40 mV and -70 mV in *Fmr1* WT and KO mice. The NMDA receptor mediated component of the EPSC was measured 60 ms after stimulation artifacts at V_m_ = +40 mV. Calibration: 20 ms, 20 pA. **B)** Grouped data from all recordings of the NMDA/AMPA ratio. **C**) Representative traces for AMPA mediated EPSCs in CA1 neurons elicited by incrementally increasing the extracellular stimulating intensities in *Fmr1* WT and KO mice. CA1 neurons were voltage-clamped at -70 mV. Calibration: 20 ms, 50 pA. **D**) Input-output curve of EPSCs in CA1 neurons. **E**) Representative traces for EPSC_NMDA_ in *Fmr1* WT and KO mice (scaled to peak). EPSCs were recorded at +40 mV in the presence of blockers of AMPA receptors and GABA_A_ receptors. Calibration: 50 ms. **F)** Grouped data for all recordings of the measured decay time constant (ι−) of the EPSC_NMDA_ **G**) Time course of EPSC_NMDA_ inhibition by GluN2B antagonist ifenprodil (IFN). Inset shows representative traces of EPSC_NMDA_ during baseline and IFN treatment (15 - 20 min bin after IFN) in WT and *Fmr1* KO slices. Calibration: 50 ms, 20 pA. **H**) Grouped data from all recordings of the % inhibition of the EPSC_NMDA_ by IFN.

### Ca2+ activated SK channel activity is upregulated by NMDA priming

The functional coupling of NMDA receptors with small conductance Ca^2+^ activated K^+^ (SK) channels has been well documented in the hippocampus where SK channels constrain synaptic NMDA receptors and regulate the induction of plasticity (Ngo-Anh et al., 2005). Activity of NMDA receptors downregulates SK channels which leads to an increase in dendritic excitability and promotes LTP induction in a positive feedback manner (Lin et al., 2008)(Lin et al., 2010)(Jones et al., 2017). Prior work has also demonstrated that SK channel currents are reduced in CA3 neurons in *Fmr1* KO mice because of a loss of a direct interaction with Fmrp (Deng et al., 2019). Therefore, we focused on these channels as potential mediators of aberrant dendritic excitability that could affect NMDA priming mechanisms. SK-mediated medium after-hyperpolarization mediating currents (I_mAHP_) were recorded in CA1 pyramidal neurons in voltage-clamp by injecting depolarizing current (Deng et al., 2019) (see methods). We found that NMDA-treated neurons exhibited larger SK-mediated currents compared to vehicle-treated neurons (F_(1.87, 97.3)_ = 3.92, p = 0.026, η²_G_ = 0.027; three-way mixed ANOVA) (Figure 5 A - D). The effect size (Cohen’s d value) of NMDA treatment was larger in *Fmr1* KO mice than in WT mice (WT: d = 0.508, 95 % CI [-0.34, 1.35]; KO: d = 0.848, 95 % CI [-0.012, 1.707]). We also recorded SK channel currents using a voltage ramp protocol (Deng et al., 2019) (see methods) and obtained similar results. NMDA treatment significantly potentiated SK currents (F_(1, 55)_ = 17.09, p < 0.001, η² = 0.23; two-way ANOVA). Post-hoc comparison showed that the effect size of NMDA treatments was large in *Fmr1* WT neurons (d = 1.00) and very large in *Fmr1* KO neurons (d = 1.16) (Figure Supplementary A - D). These results indicate that NMDA priming stimuli differentially modulates SK channel activity in *Fmr1* WT and KO mice.

**Figure 5.**
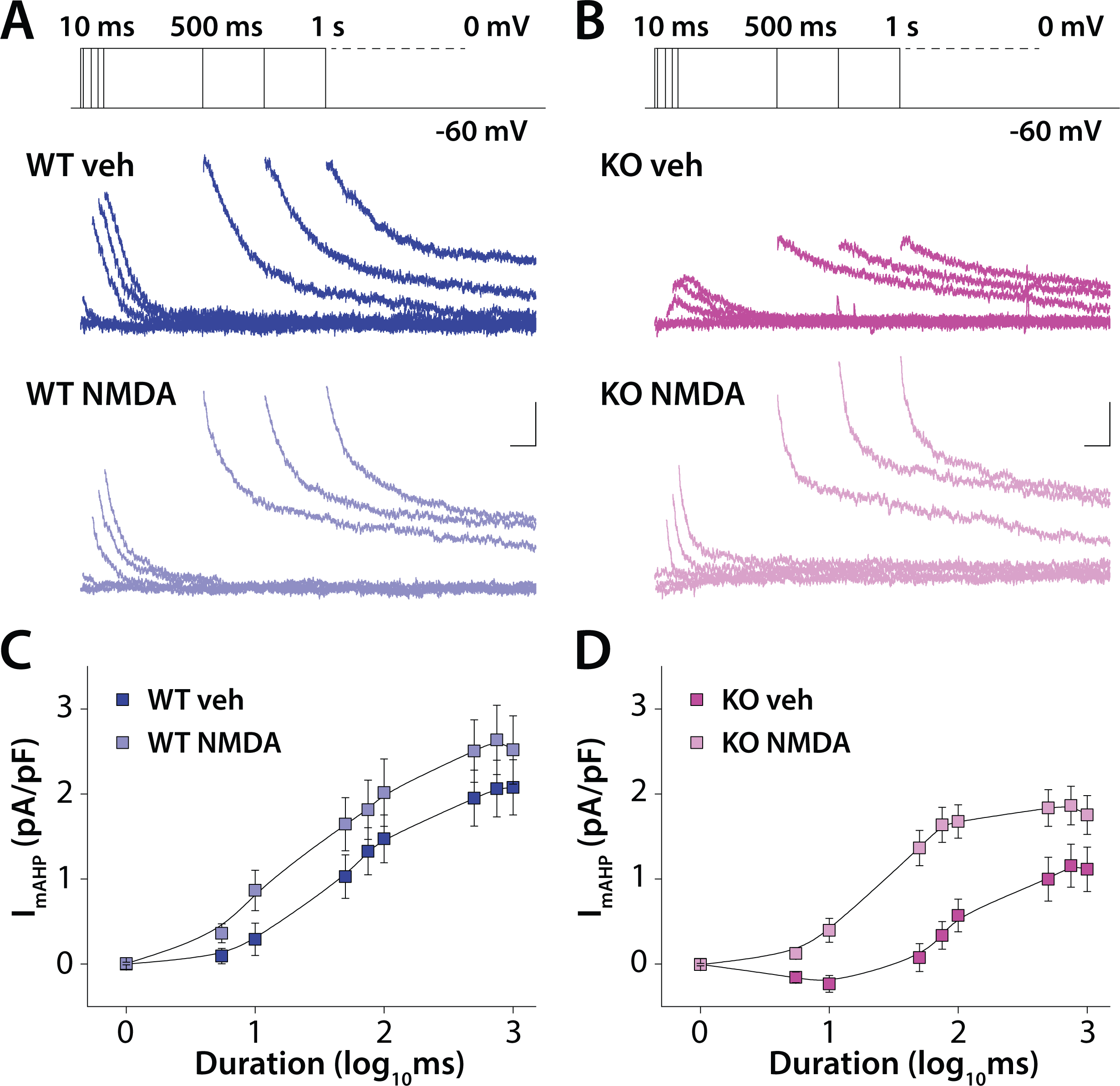
SK channel currents recorded in CA1 neurons with NMDA priming in WT and KO neurons. **A)** Top panel: voltage steps protocol from -60mV to 0mV used to activate SK currents (I_SK_) with different widths. Below: example traces at each step width in vehicle and in NMDA treatment condition for WT neuron recordings and **B**) *Fmr1* KO recordings. Calibration: 100 ms, 50 pA. **C)** Grouped data for all recordings of the I_SK_ as a function of the step duration for recordings from WT neurons in vehicle and NMDA treatment conditions. **D)** Grouped data for all recordings of the I_SK_ as a function of the step duration in *Fmr1* KO neurons in vehicle and NMDA treated conditions.

### SK channel inhibition partially restores exaggerated metaplasticity in Fmr1 KO mice

Analysis of SK channel mediated currents also detected a significant difference between *Fmr1* WT and KO mice (F_(1.87, 97.3)_ = 3.46, p = 0.038, η²_G_ = 0.024; three-way mixed ANOVA) (Figure 5 A - D). In vehicle-treated conditions, SK currents were smaller in *Fmr1* KO mice than in WT mice while it was not statistically significant (p = 0.055, d = 0.805, 95 % CI [-0.04, 1.649]) (Figure 5 A - D). These results indicate that SK channel activity is reduced in *Fmr1* KO mice in CA1 pyramidal neurons similar to previous findings in CA3 hippocampal neurons (Deng et al., 2019).

Prior work has demonstrated that acute administration of the specific SK channel blocker apamin boosts the EPSPs in CA1 neurons suggesting that SK channels normally suppress the synaptic response dampening dendritic excitability (Wang et al., 2015; Jones et al., 2017). To determine if this mechanism was active in CA1 of *Fmr1* KO mice we recorded the EPSP in CA1 and measured the effect of application of apamin (300 nM). In recordings from WT slices we found that the amplitude of the EPSP was significantly boosted by the SK channel inhibitor (WT: 131.4 ± 21.1 %, n = 7 cells, 4 mice) In contrast in recording from *Fmr1* KO animals there was no boosting of the EPSP with apamin application (KO: 82.5 ± 10.7 %, n = 7 cells, 3 mice; p = 0.024; Mann-Whitney) (Figure 6 A-C). Thus disruption of SK channels in *Fmr1* KO mice has a significant effect on synaptic potentials. The lack of effect of apamin suggests that the normal inhibitory role of SK channels in CA1 dendrites and synapses is disrupted in *Fmr1* KO mice.

**Figure 6.**
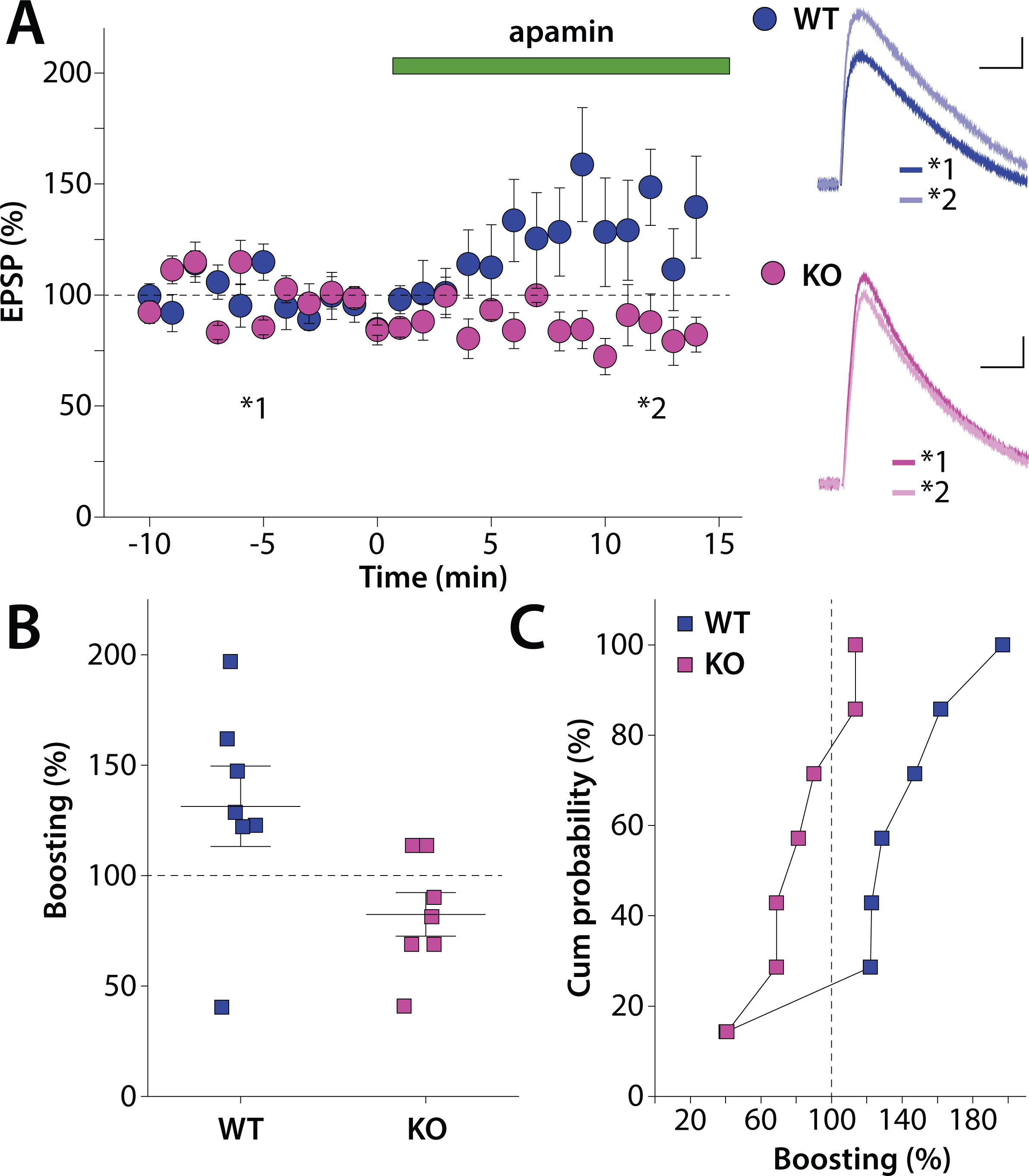
Boosting of the EPSP in CA1 by the SK channel inhibitor apamin. **A)** Time course of the EPSP with application of SK channel blocker apamin. Right inset: Representative traces of EPSPs during baseline and during apamin treatment (10 - 15 min bin after apamin treatment) from WT (top) and *Fmr1* KO (bottom). Calibration: 20ms, 1 mV. **B**) Grouped data from all recordings of the calculated % increase in the EPSP after apamin. **D**) Cumulative probability graph of apamin boosting for WT and KO recordings.

To determine whether SK channels contribute to the difference in the magnitude of NMDA mediated metaplasticity seen in *Fmr1* KO mice we measured priming induced metaplasticity in the presence of apamin (300 nM) in WT and Fmr1 KO mice. In slices from WT mice inhibiting SK channels during the 5x TBS with NMDA priming resulted in a large magnitude LTP up to 60 mins after the last TBS (WT - apamin: 171.5 ± 9.7 %, n = 10 cells, 4 mice) (Figure 7 A-C). Interestingly the magnitude of this LTP was similar to that induced in the absence of priming that we had observed previously (Figure 2 A & B) suggesting that SK had little influence on metaplasticity induced by this strong stimulus. In interleaved recordings with SK inhibition from *Fmr1* KO slices we observed equally large potentiation after NMDA priming (KO - apamin: 149.1 ± 9.8 %, n = 8 cells, 4 mice) which was not significantly different from WT recordings (p = 0.14; Mann-Whitney) (Figure 7A-C). Thus, in both these situations the magnitude of LTP was large and similar to the situation were 5x TBS LTP was induced without priming in WT mice. Taken together, our findings indicate that SK channel have an aberrant role in NMDA primed metaplasticity through a mechanism involving SK channels in *Fmr1* KO mice which is not as dominantly engaged in WT mice.

**Figure 7.**
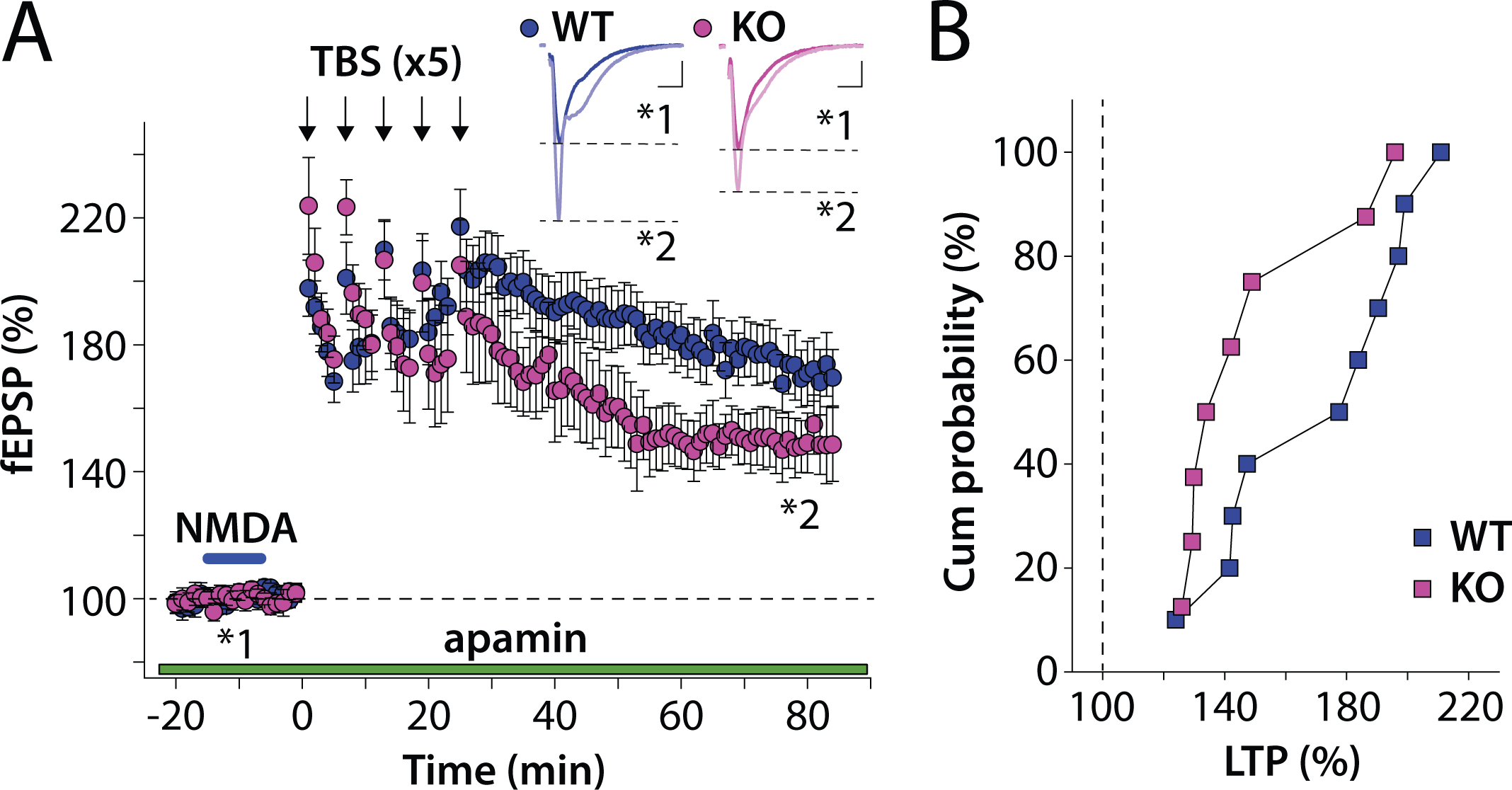
NMDA priming induced metaplasticity in the presence of apamin block of SK channels. **A)** Time course of NMDA primed metaplasticity with repeated TBS LTP in the presence of SK channel blocker apamin in WT and *Fmr1* KO slices. Inset: Representative fEPSP traces during baseline and post 5x TBS LTP condition (50 - 60 min bin after the 5^th^ TBS). Calibration: 10ms, 0.2 mV. **B**) Grouped data of LTP (%) represented in a cumulative probability distribution.

## Discussion

In this study, we found that priming with low concentrations of NMDA inhibited subsequent LTP of synapses in the CA1 subregion of the hippocampus. In WT mice, this metaplastic inhibition could be overcome by stronger LTP-inducing stimuli, but not in *Fmr1* KO mice suggesting that NMDA receptor mediated metaplasticity is aberrantly exaggerated in the absence of Fmrp. Consistent with this, NMDA priming dampened dendritic excitability to a greater extent in CA1 of *Fmr1* KO mice. We further found activity dependent SK channel upregulation was increased in the *Fmr1* KO, likely contributing to the aberrant priming. Pharmacological inhibition of SK channels restored the abnormal metaplasticity in *Fmr1* KO mice.

### NMDA-mediated metaplasticity is enhanced in Fmr1 KO mice

Metaplasticity is an activity dependent change in the state of synapses which alters their threshold for synaptic plasticity induction. Hebbian forms of plasticity such as LTP and LTD undoubtedly play central roles in brain function in both physiological and pathological conditions. However additional forms of plasticity including metaplasticity, homeostatic plasticity and intrinsic plasticity are also critical for neural circuit function (Abraham and Bear, 1996)(Turrigiano and Nelson, 2004)(Sehgal et al., 2013)(Parsons, 2018). These mechanisms interact rather than operate independently both *in vitro* and *in vivo*. For example, prior behavioral experience can induce intrinsic plasticity that then serves as metaplastic regulation of synaptic plasticity and learning (Zelcer et al., 2006)(Parsons and Davis, 2012). Our own work has linked synaptic metaplasticity *in vitro* with associative learning *in vivo* (Xu et al., 2014).

Although little is known about metaplasticity mechanism in FXS, some evidence points to alteration. In FXS individuals, transcranial magnetic stimulation (TMS) studies have reported atypical metaplasticity (Oberman et al., 2016). A TMS theta burst stimulus to primary motor cortex modulated the motor evoked potentials (MEP) and subsequent TBS enhanced the MEP more strongly in FXS individuals compared to neurotypical control subjects suggesting that a metaplastic process was elevated (Oberman et al., 2016). While these studies in humans cannot be directly mapped on *in vitro* models of plasticity, they highlight the need for detailed mechanistic investigation of metaplasticity in FXS animal models. In *Fmr1* KO mice, (S)-3,5-dihydroxyphenylglycine (DHPG) induced metaplasticity that is dependent on Grp1 mGluRs is preserved in magnitude but relies on distinct mechanisms involving protein synthesis in WT mice but is protein synthesis independent in *Fmr1* KO mice (Auerbach and Bear, 2010). Moreover a recent study demonstrated that behavioral pre-training in an oculomotor task designed to inhibit cerebellar LTD paradoxically rescued deficits in LTD-dependent learning in *Fmr1* KO mice, suggesting a lowered LTD threshold may occlude normal expression (Shakhawat et al., 2024). Our study provides additional support for common and widespread changes in metaplastic mechanisms by providing the first evidence of abnormal NMDA receptor dependent metaplasticity in *Fmr1* KO mice *in vitro*.

### SK channels contribute to exaggerated metaplasticity in Fmr1 KO mice

One possible explanation for abnormal metaplasticity in *Fmr1* KO mice is NMDAR dysfunction. However, in mice at the ages used in our study we found that the EPSC_NMDA_ was largely normal with only a small change in the kinetics which could not be put down to detectable changes in the contribution of the GluN2B subunit. Thus the aberrant metaplasticity likely involves downstream mechanisms rather than the receptors themselves.

SK channels are central regulators of NMDA receptor dependent LTP in the hippocampus (Adelman et al., 2012). SK channels undergo activity dependent downregulation following NMDA receptor activation during LTP via protein kinase A (PKA) mediated phosphorylation and endocytosis (Ngo-Anh et al., 2005)(Lin et al., 2008)(Adelman et al., 2012). However, in the present study, NMDA priming increased SK channel activity an effect that was stronger in the *Fmr1* KO mice, and likely could account for exaggerated LTP inhibition. Indeed, blocking SK channels with apamin restored LTP induction with a strong TBS after NMDA priming in KO mice to WT levels.

The mechanisms by which NMDA priming increases SK activity, particularly in *Fmr1* KO mice, remain unclear. NMDA receptor signaling pathways involved in metaplastic inhibition overlap with those mediating conventional LTD (Zorumski and Izumi, 2012) including pathways that directly affect SK channel function. For example, protein phosphatase 2A (PP2A) activates SK channels by dephosphorylating calmodulin (CaM) and increasing their open probability (Bildl et al., 2004)(Allen et al., 2007). While many signaling pathways are dysregulated in the absence of Fmrp, known mechanism that affect SK channel function are useful starting points in understanding the changes in metaplasticity associated with FXS.

### K^+^ channelopathies and dendritic dysfunction in FXS and related disorders

*Fmr1* KO mice exhibit multiple channelopathies (Brager and Johnston, 2014)(Deng et al., 2021). SK channels enriched in neuronal dendrites, regulate neuronal excitability. In this study, we identified reduced SK channel activity in *Fmr1* KO mice, consistent with prior reports (Deng et al., 2019). Dysregulation of dendritic K^+^ channels including SK channels has been implicated in FXS and other neurodevelopmental disorders (Routh et al., 2013)(Kalmbach et al., 2015)(Ordemann et al., 2021)(Nomura et al., 2023). We recently reported that SK channel downregulation contributes to dendritic hyperexcitability in a mouse model with a human variant in a glutamate receptor gene that is associated with neurodevelopmental disorder (Nomura et al., 2023). In the present study, we discovered that dendritic excitability is suppressed by NMDA priming, which was likely due to an increase in SK channel activity. Given our and other studies findings it is possible that dendritic dysfunction caused by K^+^ channelopathies is a convergent endophenotype in multiple forms of neurodevelopmental disorders.

In conclusion, our findings suggest that synapses in *Fmr1* KO mice are more vulnerable to prior activity leading to maladaptive tuning of plasticity thresholds. NMDA receptor dependent metaplasticity driven by activity dependent regulation of dendritic K^+^ channels is aberrantly enhanced in *Fmr1* KO mice. We propose that pathological coupling between NMDARs and SK channels represents a key synaptic mechanism in FXS. Although the precise regulatory pathways remain to be determined, targeting NMDA receptors and SK channels may provide a therapeutic strategy to restore proper synaptic plasticity in FXS.

## Materials and Methods

### Animals

All experimental procedures were conducted in accordance with the ethical policies and protocols approved by the Northwestern University Institutional Animal Care and Use Committee (IACUC). *Fmr1* KO mice (C57/bl6J strain) were purchased from Jackson laboratory (JAX #003025). Heterozygous KO (*Fmr1^+/-^*) female mice were crossed with *Fmr1* WT (*Fmr1^+/y^*) or KO (*Fmr1^-/y^*) male mice for breeding, and male *Fmr1* WT and KO littermates (3 – 5 weeks old) were used for experiments. Mice were housed with food and water provided *ad libitum* under 12/12 h dark-light cycle. Experiments were conducted with the investigator blind to the genotype of the animals. These were subsequently confirmed by *post hoc* PCR using tail biopsy samples.

### Electrophysiology

Mice were deeply anesthetized with inhalation of isoflurane and an intraperitoneal injection of xylazine (10 mg/kg) and ketamine (100 mg/kg). Mice were perfused transcardially with an ice-cold sucrose ACSF solution containing the following (in mM): 85 NaCl, 2.5 KCl, 1.25 NaH_2_PO_4_, 25 NaHCO_3_, 25 glucose, 75 sucrose, 0.5 CaCl_2_, and 4 MgCl_2_, including 10 μM DL-APV and 100 μM kynurenate. Horizontal brain sections (350 µm thick) were prepared in the same ice-cold sucrose ACSF on a Leica Vibratome (Leica Microsystems). Slices were incubated in sucrose ACSF at 30°C for ∼30 min, which was gradually exchanged for a recovery ACSF containing the following (in mM): 125 NaCl, 2.4 KCl, 1.2 NaH_2_PO_4_, 25 NaHCO_3_, 25 glucose, 1 CaCl_2_, and 2 MgCl_2_, with 10 μM DL-APV and 100 μM kynurenate at room temperature (RT). For recordings, slices were transferred to a recording chamber after a recovery period of at least 1.5 h. Slices were visualized under Dodt-Gradient-Contrast optics (Luigs & Neumann). Recording chamber was continuously perfused with ACSF (28°C-30°C) containing the following (in mM): 125 NaCl, 2.4 KCl, 1.2 NaH_2_PO_4_, 25 NaHCO_3_, 25 glucose, 2 CaCl_2_, and 1 MgCl_2_ equilibrated with 95% O_2_ and 5% CO_2_. Standard techniques for extracellular field or patch clamp recordings were applied for electrophysiological recordings from hippocampal CA1 pyramidal neurons. For voltage clamp recordings electrodes had tip resistances of 3-5 MΩ when filled with a cesium-based internal solution containing the following (in mM): 95 CsF, 25 CsCl, 10 Cs-HEPES, 10 Cs-EGTA, 2 NaCl, 2 Mg-ATP, 10 QX-314, 5 tetraethylammonium (TEA)-Cl, and 5 4-AP. For current clamp recordings a potassium-based internal solution was used containing the following (in mM): 125 KMeSO_4_, 5 KCl, 5 NaCl, 0.02 EGTA, 11 HEPES, 1 MgCl_2_, 10 phosphocreatine, 4 Mg-ATP, 0.3 Na-GTP. Liquid junction potential was not corrected. In voltage-clamp recordings, access resistance (R_a_) was continuously monitored, and data were excluded when R_a_ showed >20% change during the experiments. Neurons were voltage-clamped at −70 mV unless otherwise specified. For extracellular recordings, normal ACSF same as the recording extracellular solution was used in the pipette. Data were acquired and analyzed using a Multiclamp 700B amplifier and pClamp 10 software (Molecular Devices).

Monopolar glass electrode filled with normal ACSF were placed in the *stratum radiatum* to stimulate synaptic afferents from CA3 pyramidal neurons (Schaffer collaterals). EPSCs were recorded in the presence of GABA_A_ receptor antagonist picrotoxin (PTX) (50 μM). The ratio of EPSC_NMDA_ to EPSC_AMPA_ was measured by switching the holding voltage from -70 mV to +40 mV, in which the measured amplitude of the EPSC at +40 mV holding 60 ms after the stimulation was considered to be mediated by NMDA receptors. NMDA receptor mediated EPSCs (EPSC_NMDA_) were isolated by an inclusion of AMPA and kainate receptor blockers CNQX (10 μM) into the extracellular recording solution and were recorded at +40 mV holding potential. The kinetics of EPSC_NMDA_ was determined by fitting the currents to a single exponential equation *y = Ae^-x/−^ + b* from which the time constant tau (τ) was derived. NMDA receptor subunit composition was estimated by measuring ifenprodil (IFN) (5 μM) mediated inhibition of the EPSC_NMDA_.

LTP was recorded in extracellular recording configuration. Stimulating and recording electrodes were placed in the *stratum radiatum* to record field EPSPs (fEPSPs) in the CA3 – CA1 synapses. LTP was induced by theta burst stimulation (TBS) in which a 5- pulse burst at 100 Hz was delivered 15 times at 5 Hz. TBS was applied either once or 5 times with 2 min inter-TBS intervals as LTP inducing stimuli. NMDA mediated metaplasticity was examined by a bath application of a low concentration of NMDA (1 μM) for 10 min in the extracellular recording solution before TBS application. To analyze the effect of SK channels, a selective blocker apamin (300 nM) was bath applied in the extracellular recording solutions throughout recordings.

SK channel activity was measured in voltage-clamp configuration using a K^+^-based internal pipette solution. Tetraethylammonium (TEA) (1 mM) and tetrodotoxin (TTX) (1 μM) were included in the external recording solutions to block other K^+^ channels and Na^+^ channels, respectively. Currents were recorded in the absence and presence of the SK channel blocker apamin. The currents recorded in apamin treated condition were then subtracted from the currents recorded in baseline condition to isolate the apamin sensitive SK channel mediated currents. Medium AHP mediating currents (I_mAHP_) were evoked by depolarizing cells from −60 to 0 mV with multiple depolarization steps with different step durations (from 1 to 1000 ms). Currents were measured 20 ms after the end of the step so as not to record contaminated capacitance transients. Ramp evoked currents were recorded by depolarizing cells from -100 mV to -25 mV (5 mV/s; 15 s). Currents were averaged from a 20 ms interval and compared at Δt = 15 s bins (corresponding to V_m_ = -24.99 mV to -25 mV bin) between groups (Deng et al., 2019). SK channel mediated currents were compared between vehicle (ACSF) treated or NMDA (1 μM) treated groups in *Fmr1* WT and KO mice.

### Ca^2+^ imaging

Two-photon laser scanning microscopy (2PLSM) was used to image CA1 pyramidal neurons in slices using methods similar to those previously described (Nomura et al., 2023). Cells were filled with AlexaFluor-568 (50 μM) through the recording electrodes for visualization. Neurons were perfused with the dye for at least 20 min after forming the whole-cell configuration. Images were acquired using femtosecond pulsed laser excitation at 790 nm (Mira 900P with a Verdi 10W pump, Coherent Laser). Laser power was controlled with an M350 (KD*P) series Pockels cells (ConOptics). A Prairie Ultima (Bruker Nano, Fluorescence Microscopy Unit) scan head on an Olympus BX-61 upright microscope was used for imaging the slice with two Hamamatsu R3982 side on photomultiplier tubes. For Ca^2+^ imaging, cells were loaded with the Ca^2+^ indicator Fluo-4 (200 μM) through the recording patch electrodes. Dendritic Ca^2+^ signals were triggered by 10 back-propagating action potentials (bAPs) evoked by somatic suprathreshold depolarizing currents (1 nA for 1 ms at 100 Hz). ROIs were localized in the apical dendrites of CA1 pyramidal neurons ∼100 μm distal from the soma. Ca^2+^ signals were acquired as fluorescent line scan signals in 3-5 pixels (0.77 µm) per line with 22.8 µs dwell time. To analyze the effect of NMDA priming, slices were treated with a low concentration of NMDA (1 μM) for 10 min as in electrophysiological recordings. Ca^2+^ signals were compared in the same ROIs in the same cells between baseline and NMDA treated conditions. Recordings were configured and acquired using a Multiclamp 700B amplifier and Prairie View 9.0 software (Bruker).

### Experimental design and statistical analyses

Experimental design, statistical tests, and critical statistical values in each experiment are presented in the main text. Excel (Microsoft), GraphPad Prism (GraphPad Software), JASP (U Amsterdam), and Origin (OriginLab) software were used for statistical analysis. Comparisons were made using Mann–Whitney *U* test or Kolmogorov–Smirnov test for unpaired two samples. Paired two samples were compared using Wilcoxon signed-rank test. For multiple comparisons, Kruskal-Wallis test, two-way ANOVA, repeated two-way ANOVA, or three-way mixed-design ANOVA with one repeated measure factor followed by *post hoc* Bonferroni’s correction were used. Sphericity was tested by Mauchly’s test. Greenhouse–Geisser-adjusted degrees of freedom were used when appropriate. Differences were considered to be significant when *p* < 0.05. Data are presented as mean ± SEM. This study was not preregistered. All data presented in the study will be available upon requests.

## Author Contribution

T.N. and A.C. designed research, T.N. and C.M. performed research, T.N., C.M., and A.C. analyzed data, T.N, C.M., and A.C. wrote the paper.

## Conflict of interest

The authors declare no competing financial interests.

## Acknowledgements

This work was supported by NIH/NIMH R01MH099114 and NIH/NICHD R01HD108370 to AC. TN was supported by Research Grant from National Center of Neurology and Psychiatry (NCNP) (Japan). CM was supported by R25 GM121231 from NIGMS. The authors thank members of Contractor lab for helpful inputs to the manuscripts in particular Drs. John Marshall and Jian Xu for help with statistical analyses.

## Figure Legends

**Figure S1.**
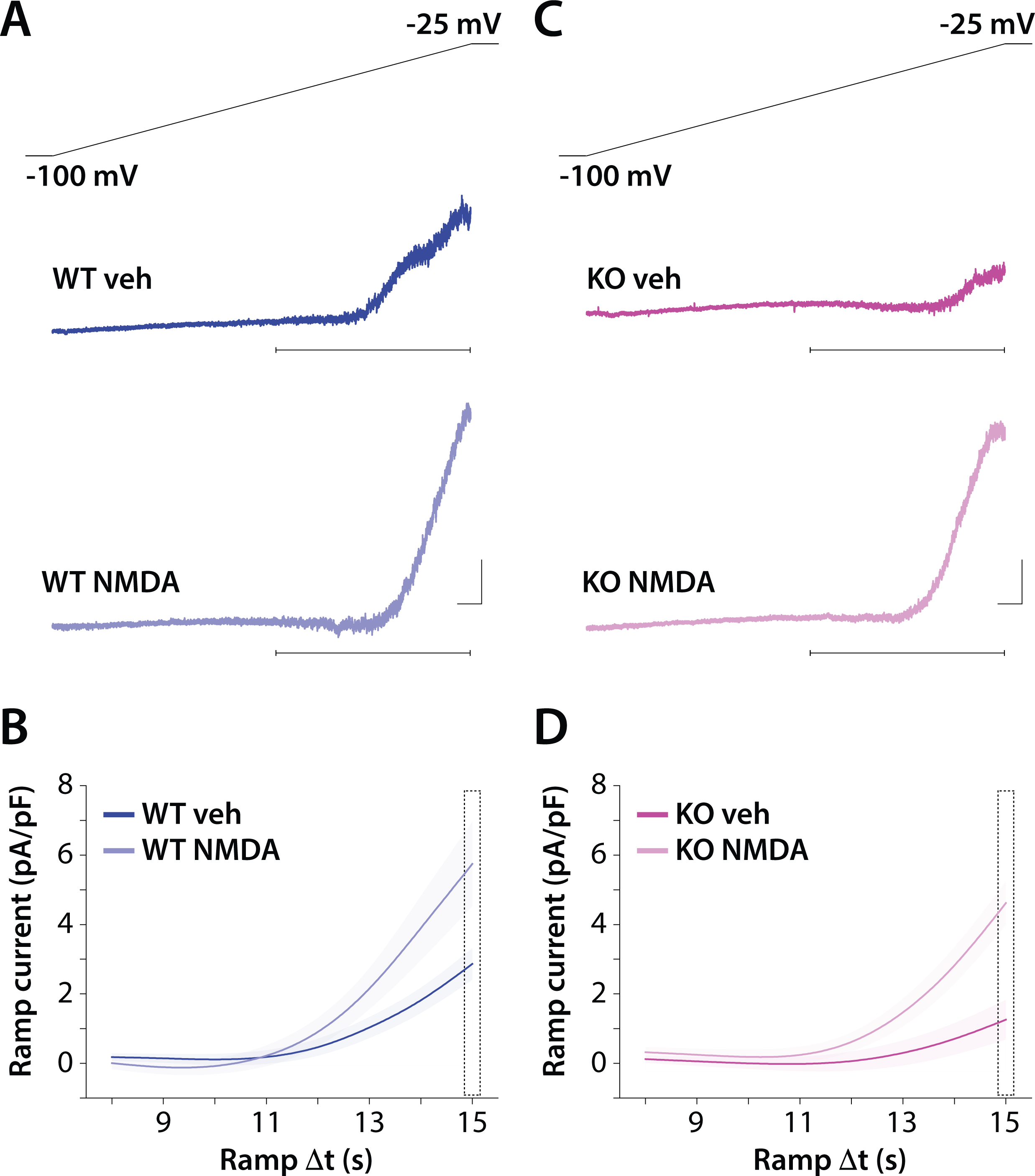
SK channel activity recorded using a voltage ramp protocol. **A)** Top shows ramp protocol and lower panels are current traces in vehicle treated condition and NMDA treatment condition for WT recordings. Calibration: 1 s, 100 pA. **B**) Average current of ramp Λ1t of 8 – 15 s (indicated by horizontal bars in panel **A**) as a function of time along the ramp for WT recordings in vehicle and NMDA treatment conditions. **C**) Ramp protocol and example current traces recorded in KO neurons. Calibration: 1 s, 100 pA. **D**) Average current as a function of ramp time for recordings from *Fmr1* KO neurons in vehicle and NMDA treated condition.

## Notes

### Competing Interest Statement

The authors have declared no competing interest.

